# Cryptochrome Stabilization Ameliorates Chronic Pain

**DOI:** 10.1101/2022.07.15.499708

**Authors:** L. Wei, Y. Wu, F. Zirpel, G. Flower, J. V Vierbergen, V. Wieser, S. Peirson, H. Sleven, A. Pokhilko, M. Z. Cader

**Affiliations:** Nuffield Department of Clinical Neurosciences, University of Oxford, Dorothy Crowfoot Hodgkin Building, South Parks Road, Oxford, OX1 UK

## Abstract

Physiological and pathological pain exhibits striking diurnal variation, but the underlying mechanisms are largely unknown. We now describe an independent molecular clock in peripheral sensory neurons and satellite glial cells of sensory ganglia. We show that it is the sensory neuron transcription-translation feedback loops (TTFLs) that are responsible for diurnal pain behaviors. This clock regulates diurnal neurophysiological responses to a range of ligands, as well as synaptic activities of primary nociceptors. Furthermore, we find that loss of *Cry1* and *Cry2*, the repressive arm of the core TTFLs, intensifies pain responses associated with increased voltage-gated sodium channel currents. Conversely, stabilization of CRY1 and CRY2 using the small molecule KL001, reduces pain sensitivity. Our results highlight novel opportunities to address chronic pain by directly harnessing circadian mechanisms.

**One-Sentence Summary:** A peripheral pain clock governs daily pain fluctuations, which can be harnessed for treating pain disorders.

---

Pain is an essential aversive sensory experience that enables awareness of tissue injury or threat of tissue damage, and as such is critical for survival. Pain sensation, nociception, is mediated by peripheral sensory neurons, which express ion channels and receptors, transducing noxious stimuli into action potentials and gating synaptic transmission of nociceptive information in the spinal cord and lower brainstem. The repertoire of ion channels and associated signaling networks, as well as synaptic proteins, is generated by the cell bodies of afferent sensory neurons residing in sensory ganglia, ensheathed by satellite glial cells (SGC). Nerve injury, primary dysfunction of pain pathways and inflammatory activation of peripheral nociceptors can lead to a range of chronic pain disorders. A characteristic feature of chronic pain is diurnal variation in pain severity, with different conditions showing peaks at particular times – for example, migraine peaks in the early morning(*1*), carpal tunnel syndrome exacerbates at night(*2*) and trigeminal neuralgia is typically quiescent at night(*3*). Treatments for chronic pain remain inadequate and the fluctuations in pain severity can be particularly challenging for patients, disrupting sleep and daily activities.

It has been long established that pain sensory thresholds exhibit pronounced diurnal variation(*4, 5*) but the mechanisms responsible for such variation is unknown. Diurnal patterns can arise for a variety of reasons, particularly through entrainment to light-dark cycles. However, intrinsic circadian behaviors will continue under constant conditions. We, therefore, investigated whether pain rhythmicity observed in 12:12 Light-Dark (LD) was retained in constant darkness (DD). As expected, we found diurnal variation in punctate mechanical sensitivity in the hindpaw as assessed by von Frey filaments and orofacial nociceptive responses to a von Frey filament (Supplementary Fig. 1A-B). The 50% withdrawal threshold is at the lowest level in the first part of the light period whilst the highest tolerance is during the early hours of darkness (ZT12-15h), before dropping again (ZT18-21h). Similarly, the orofacial nociceptive response to mechanical stimulation with a 0.4g von Frey filament followed the same variation in responses, with significantly heightened nociception scores during the early day and reaching a nadir at ZT12-15. We found that female wildtype mice exhibited the same patterns as males (Supplementary Fig. 2). The diurnal pattern was preserved under constant DD conditions, for both von Frey filament responses of the hindpaw and orofacial nociceptive responses (Supplementary Fig. 1). The variation in pain sensitivity was highly consistent over several days of repeated testing (Fig. 1A-B). We next assessed if this circadian pattern persisted in a chronic pain model, using intraperitoneal (i.p.) injections of nitroglycerin (GTN), which is a well-established model of migraine(*6-8*). Mice were administered GTN (10mg/kg) on alternate days with behavior assessed the day after injection for 10 days in total. As expected, responses to von Frey filaments and plantar thermal sensitivity measured using the Hargreaves method were exaggerated compared to vehicle control after repeated GTN, indicating peripheral sensitization for mechanical and thermal responses. Importantly, circadian variation of sensitized responses was present in mice injected with GTN (Fig. 1A-B, Supplementary Fig. 1). This indicates that the mechanisms driving the circadian variation in nociceptive responses are preserved even in chronic pain states.

**Fig. 1:**
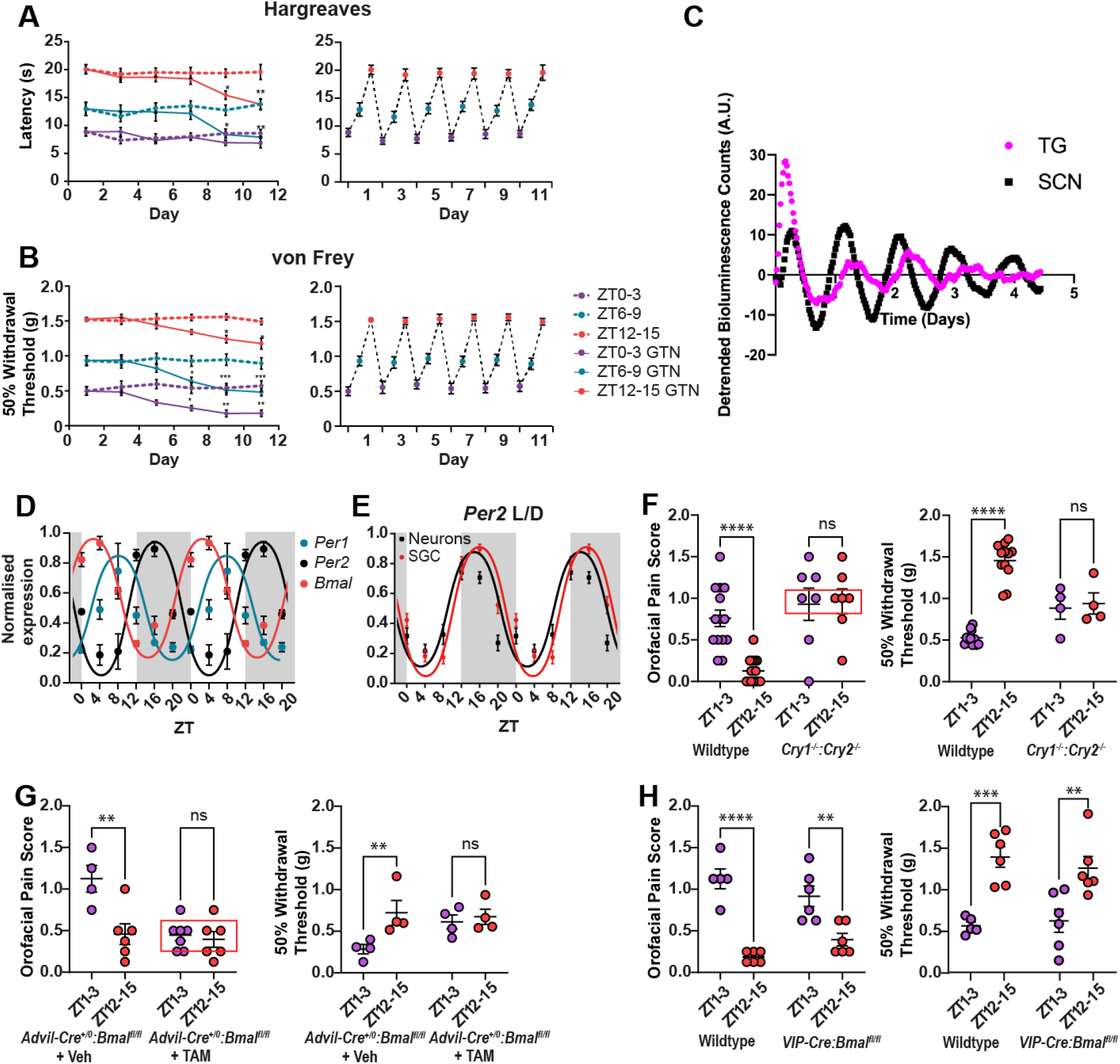
An autonomous peripheral sensory neuron clock regulates diurnal pain responses. Hargreaves test (**A**) and von Frey hindpaw test (**B**) showed steady and persistent differences in thresholds between the three time windows tested (ZT 0-3; ZT 6-9; ZT 12-15) over 11 days both at baseline and after chronic i.p. 10mg/kg GTN administration. 3 wildtype mice per data point, n=18. Two-way ANOVA, significant interaction between days and treatment (**A**) F_(25,60)=_2.272, p=0.005 and (**B**) F_(25,60)_=2.839, p=0.0005. (**C**) Bioluminescence of explanted TG and SCN tissue from *Per2-luciferase* reporter mouse over 5 days. The graph shows representative data from one mouse. (**D**) Gene expression of three clock genes (*Per1, Per2, Bmal1*) at 6 time points. 3 wildtype mice per time point, n=18. Data fitted to sine wave. (**E**) *Per2* gene expression in SGC and TG neurons isolated from 3 wildtype mice per time point, n=18. Data fitted to sine wave. (**F-H**) von Frey mechanical hindpaw thresholds and orofacial pain response between two time points (ZT1-3 or ZT12-15). (**F**) Wildtype (n=14-15 per time point) and *Cry1*^-/-^:*Cry2*^-/-^ mice (n=7). Two-way ANOVA significance between time and genotype F_(1,19)_=16.6, p=0.0006 for orofacial and F_(1,27)_=29.8, p=<0.0001 for hindpaw. Red box highlights mean orofacial response of *Cry1*^-/-^:*Cry2*^-/-^ mice. (**G**) *Advillin-Cre-ERT2:Bmal* ^*fl/fl*^ mice induced with tamoxifen (TAM) (n=6-7 per time point) or vehicle (n=4-6 per time point). Two-way ANOVA showed significant interaction between time and genotype F(_1,19)_=8.16, p=0.01 for orofacial and F_(1,6)_=8.558, p=0.0264 for hindpaw. Red box highlights mean orofacial response of sensory neuron *Bmal1* knocked out mice. (**H**) Wildtype (n=5-6 per time point) and *VIP-Cre:Bmal* ^*fl/fl*^ mice (n=6 per time point). Two-way ANOVA showed a significant interaction between time and genotype of orofacial F_(1,19)_=5.055, p=0.0366. For hindpaw, significance was observed for time F_(1,19)_=34.38, p<0.0001. (**F-H**) All grouped data are mean +/- SEM. * p<0.05, ** p<0.005, *** p<0.0005, **** p<0.0001

Peripheral sensation is transduced by primary sensory neuronal afferents that arise from either the dorsal root ganglia (DRG) or trigeminal ganglia (TG). We sought to investigate whether these structures may have autonomous clocks. We utilized *Per2-Luciferase(Luc)* knock-in mice to monitor the bioluminescence associated with real-time *Per2* expression. TG and suprachiasmatic nuclei (SCN) were dissected from PER2::LUC mice kept in LD conditions and bioluminescence was recorded every 30 minutes for approximately 5 days with no media change (Fig. 1C). Consistent with previous literature, the SCN maintains a robust and sustained circadian *Per2* rhythm throughout the recording. In the TG, circadian *Per2* oscillations are also clearly observed. Both tissues have a similar period but TG circadian oscillations were delayed relative to the SCN peak by approximately 3h. In order to confirm clock gene oscillations *in vivo*, RT-qPCR of *Per2, Per1*, and *Bmal1* was performed on TG samples dissected every 4h during one day for 6 time points in total (ZT: 0, 4, 8, 12, 16, 20) and clear oscillations were observed (Fig. 1D). This aligns with the recent identification of a peripheral clock in DRG sensory neurons(*9*)

The TG is composed of sensory neuron cell bodies ensheathed by SGCs as well as Schwann cells that myelinate the larger fibers. We sought to understand from which cell type (neuron or glia) the circadian rhythms were originating. After dissection of TGs at 6 time points in LD cycle, cells were dissociated into glial and neuronal fractions by density-gradient centrifugation. We confirmed the successful enrichment of each fraction by expression of glial and neuronal markers *Gfap* and *Trpv1*, using RT-qPCR (Supplementary Fig. 3); these markers do not show circadian oscillations. We found that *Per2* expression was oscillating in both glia and neurons, in close phase to each other (Fig. 1E).

Molecular clocks are present in many tissues and we first sought to establish the dependence of pain behaviors on the transcription-translation feedback loops (TTFLs) by using *Cry1*^-/-^:*Cry2*^-/-^, that lack the critical repressive arm of the clock(*10*). We found that with global deletion of the clock, mice lost their variation in sensory responses (Fig. 1F). To more specifically examine whether the local clock in peripheral sensory neurons might drive the observed behaviors, we used an *Advillin-Cre-ERT2* mouse crossed with a floxed *Bmal1* mouse(*11*). Following tamoxifen injection, inducing *Bmal1* deletion only in the sensory neurons (Supplementary Fig. 4), we found that the diurnal differences in pain responses were no longer present (Fig. 1G). In contrast, when we ablated the SCN clock through floxed *Bmal1* deletion in *VIP-Cre* mice (*12*), we found diurnal pain behaviors were retained (Fig. 1H). This suggests that whilst TTFLs in SCN VIP neurons are not required, the TTFL in DRG and TG primary sensory neurons have autonomous TTFLs and are required for pain circadian rhythms.

Our results demonstrate that during the light-on period (ZT4) the animals display greater pain sensitivity and responses and display the lowest pain sensitivity and responses at the start of the lights-off period (ZT16), during their active phase. This coincides with the *Per2* trough (ZT4) and *Per2* peak (ZT16). We therefore next sought to assess how clock genes might lead to variation in pain behaviors by investigating neurophysiological responses of primary nociceptive afferents. We examined neuronal and glial responses in intact acute TG slices, using Fura2-AM ratiometric calcium imaging upon exposure to capsaicin, menthol, or ATPγS. In wildtype animals the responses to these ligands displayed clear diurnal patterns, with a greater proportion of cells responding at ZT4 compared to ZT16 (Fig. 2A). In *Cry1*^-/-^:*Cry2*^-/-^mice, these diurnal patterns were disrupted in both neurons and satellite glia (Fig. 2B). Using multi-electrode arrays (MEA), we recorded spontaneous firing from TG slices and, unexpectedly, we found that spontaneous/basal activity was greater at ZT16 (Fig. 2C). However, application of ATPγS led to significantly more neuronal activity only at ZT4, but not ZT16. We also investigated the central synaptic activity of primary nociceptors by performing patch clamp recordings from cell bodies from layers I/II of acute slice of the trigeminal nucleus caudalis (TNC). We found that the frequency of excitatory postsynaptic currents (sEPSC) was significantly increased at ZT16 compared to ZT4 in wildtype animals and this pattern was lost in *Cry1*^-/-^:*Cry2*^-/-^ mice (Fig. 2D) – consistent with results from the MEA assays.

**Fig. 2:**
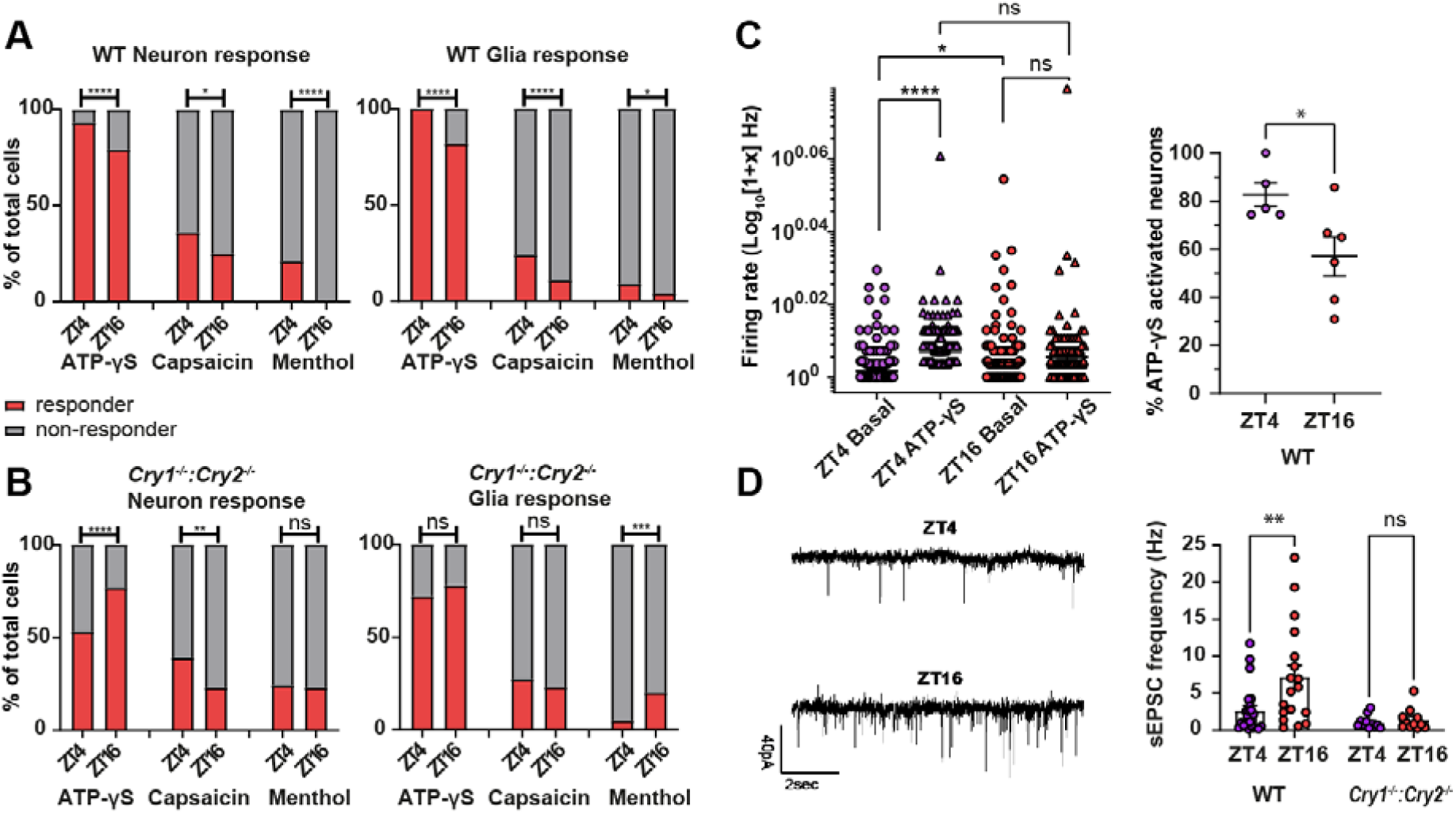
Circadian electrophysiological properties of trigeminal ganglia (TG). (**A-B**) Fura2-AM ratiometric calcium imaging showed diurnal differences in the proportion of responsive neurons and glia cells to ATP-□S 50uM, capsaicin 10uM, menthol 200uM at ZT4 and ZT16 of wildtype mice **(A)** or *Cry1*^-/-^: *Cry2*^-/-^mice (**B**) TG slices. Chi-square test with two-tail, * p<0.05, ** p<0.005, *** p<0.0005, **** p<0.0001 (**C, left panel**) Firing rate of primary mouse TG neurons, before and after stimulation with ATP-□S 50uM, recorded from acute TG slices using multiple electrode arrays (MEA) at ZT4 and ZT16. Data log normalized. One-way ANOVA *p=0.0111, ***p=0.0001, ****p<0.0001. (**C, right panel**) MEA electrode activation after addition of ATP-□S 50uM per TG slice between ZT4 and ZT16 (TG slices n=11). Unpaired two-tail t-test, *p=0.0308. (**D**) Patch clamp current traces **(left panel)** and frequency **(right panel)** of spontaneous excitatory postsynaptic current (sEPSC) in Lamina I/II of trigeminal nucleus caudalis neurons from acute slices of wildtype mice or *Cry1*^-/-^:*Cry2*^-/-^mice at ZT4 and ZT16. 2-way ANOVA showed significant interaction between time and genotype F_(2,81)_=3.226, p=0.0449, and significance between time points in wildtype **p=0.0025.

Neurophysiological responses of primary sensory neurons are regulated by the expression of voltage and ligand-gated ion channels, particularly potassium and sodium channels. We performed RT-qPCR on a candidate set of genes on TG dissected from animals over 6 time-points in 24 hours. The genes encoding receptors for nociceptive ligands such as *Trpv1, Trpa1, Trpm8, P2×4*, and *P2×5* did not show significant diurnal variation (Supplementary Fig. 5A). This suggests the observed diurnal variation in calcium responses to capsaicin, ATPγS and menthol (Fig. 2A) is not due to circadian regulation of transcription of these genes. However, the expression of *Kcnk2, Kcnk10, Kcnk18, Scn8a, Scn9a, Nr1d2, Nr1d1*, and *Kcna1* showed significant diurnal expression (Supplementary Fig. 5A-B). This pattern was lost in the *Cry1*^-/-^ :*Cry2*^-/-^ mice, confirming the role of the TTFLs in driving the oscillations of these ion channel genes.

Expressed ion channel gene transcripts are translated, assembled, and trafficked to the membrane and it is, therefore, essential to investigate whether ionic currents correspond to diurnal gene expression patterns. We used primary TG cultures, synchronized with forskolin (Supplementary Fig. 6A-D) to investigate small diameter nociceptor properties and ionic currents at a time when *Per2* is low and *Per2* is high, matching the time points when mice show increased or lowered pain sensitivity respectively. Strikingly, the voltage dependent sodium currents were much larger at *Per2-*low time-point in comparison to *Per2-*high time-point. In *Cry1*^-/-^:*Cry2*^-/-^ mice, the diurnal variation was lost, and instead, these animals displayed increased sodium currents at both time points (Fig. 3A-B). We found that resting membrane potential did not show diurnal variation, in contrast to tissues such as the SCN(*13*), whilst rheobase was slightly elevated (Fig. 3D, F). This may correspond to the increase in spontaneous neuronal activity observed at ZT16 compared to ZT4 (Fig. 2C, D). The leak potassium current between the two time points was not significantly different (Fig. 3E).

**Fig. 3:**
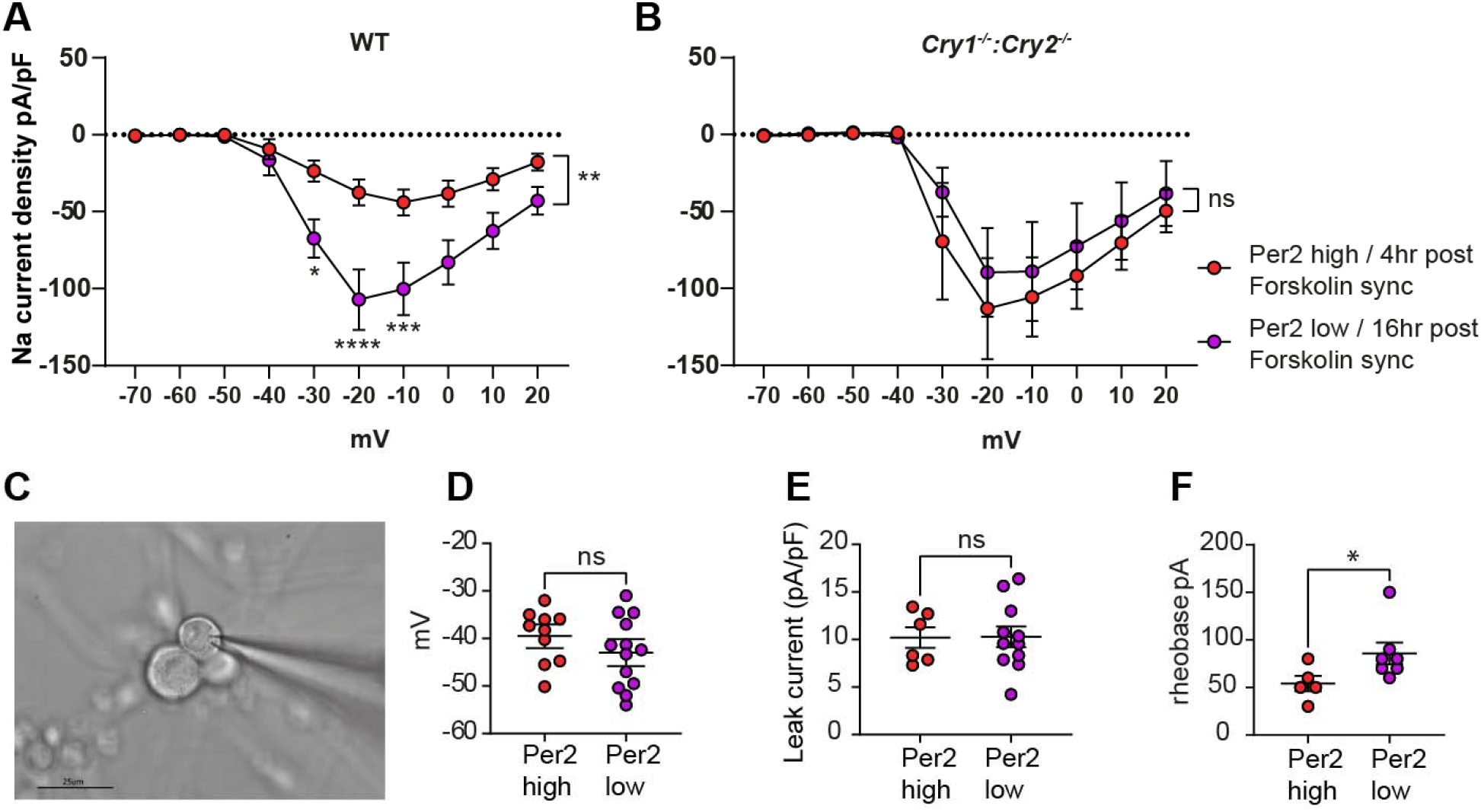
Clock dependent primary nociceptor neuronal properties. (**A-B**) patch clamp recordings of voltage dependent sodium currents in wildtype (n=11 per time point) **(A)** or *Cry1*^-/-^ : *Cry2*^-/-^ (n=7 per time point) **(B)** mouse primary nociceptors, following 10µM forskolin synchronization, at time points where per2 were low or high. 2-way ANOVA showed significant interaction between time and current for wildtype F_(9,180)_=5.081, p=<0.0001, and no significant interaction for *Cry1*^-/-^: *Cry2*^-/-^ F(9,108)=0.303, p=0.9724. (**C**) Brightfield image of primary nociceptor and attached patch electrode. (**D-F**) intrinsic neuronal properties in wildtype mouse primary nociceptors at per2 low and per2 high time points following 10µM forskolin synchronization - resting membrane potential (n=23 per time point) **(D)**, potassium leak current (n=17 per time point) **(E)**, and rheobase (n=12) **(F)**. Unpaired two-tail t-test. * p≤0.05.

CRY1 and 2 are transcriptional repressors of the clock and we noticed that in comparison to the wildtype responses, *Cry1*^-/-^: *Cry2*^-/-^ mice displayed increased pain sensitivity at ZT16 (Fig. 1F). BMAL1 is a transcriptional activator of the clock, inducing gene expression, and in *Advillin-Cre:Bmal*^-/-^ pain sensitivity at ZT4 is reduced (Fig. 1G). We, therefore, hypothesized that stabilization of CRY1/CRY2 would reverse the increased voltage dependent sodium currents and thus reduce pain sensitivity. We used KL001 which was previously identified from a small molecule screen of CRY1/CRY2 stabilizers (*14*). The application of KL001 to forskolin synchronized primary TG cultures resulted in a significant decrease of the sodium current (Fig. 4A). We then administered KL001 or vehicle control intraperitoneally (i.p.) to wildtype mice at ZT0 and investigated the effect at ZT4 when pain sensitivity was increased. We found that KL001 substantially decreased pain sensitivity in both hindpaw to von Frey stimulation, as well as orofacial nociceptive responses (Fig. 4B). This decrease in sensitivity was not observed in the *Cry1*^*-/-*^*:Cry2*^*-/-*^ mice injected with KL001, confirming the specificity of KL001 effects on pain through CRY1 and CRY2 (Supplementary Fig. 7). We next administered KL001 or vehicle to mice two hours before acute i.p. GTN injection and found that mice injected with KL001 had significant pain reduction compared to the control group. Finally, we repeated this experiment in the chronic GTN model, where KL001 but not vehicle significantly reduced chronic orofacial pain sensitization induced by repeated GTN injections (Fig. 4C).

**Figure 4:**
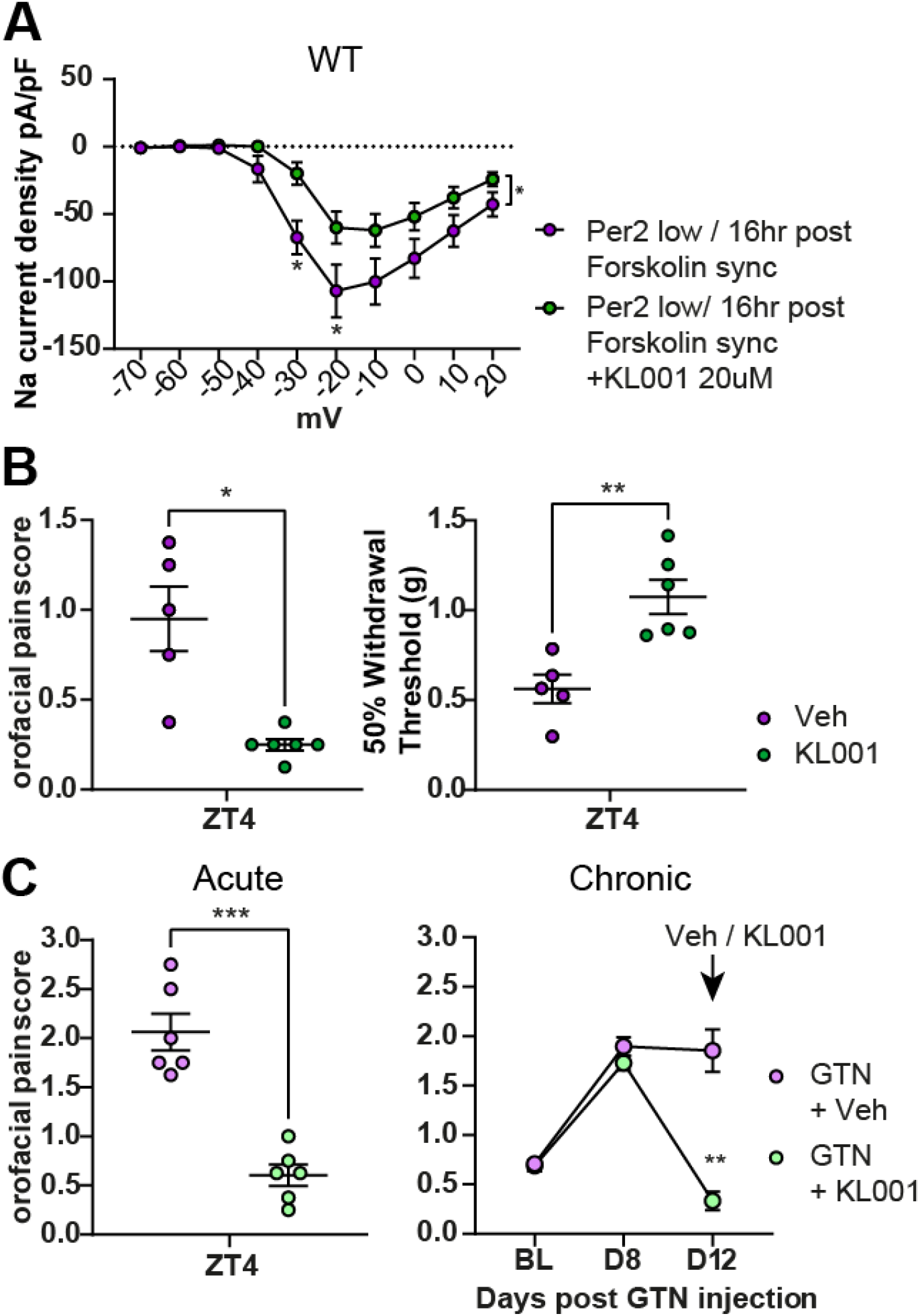
The small molecule CRY1/2 stabilizer (KL001) decreases voltage dependent sodium currents and pain responses. (**A**) patch clamp recordings of voltage dependent sodium currents in wildtype mouse primary nociceptors, following 10µM forskolin synchronisation at per2 low time point with or without 20µM KL001. 2-way ANOVA showed significant interaction between time and current F_(9,153)_=2.106, p=0.0322 (vehicle n=11, KL001 n=8). (**B**) orofacial pain response and von Frey mechanical hindpaw thresholds at ZT4 in wildtype mice i.p. injected with vehicle (n=5) or 0.4mg KL001 (n=6); unpaired two-tail t-test * p≤0.05, ** p<0.005. (**C**) orofacial pain responses in wildtype mice administered vehicle (n=6) or 0.4mg KL001 (n=6) in acute (left) or chronic GTN model from baseline (BL) to day 12 (D12) (right); unpaired two-tail t-test *** p<0.0005 and two-way ANOVA ** p=0.0011. All grouped data are mean +/-SEM, * p<0.05, ** p<0.005, *** p<0.0005, **** p<0.0001.

Collectively our data has revealed the importance of the peripheral sensory neuron clock in determining circadian pain responses. Furthermore, we have found the clock establishes diurnal differences in neurophysiological properties, including large differences in currents mediated by voltage gated sodium channels. Cryptochromes, Cry1 and 2, are important gene repressors, and our data suggests that their stabilization can engage the downstream effector mechanism that reduces pain responses. There are a growing number of medicines targeting the molecular clock, and these represent novel means to treat chronic pain.

## Acknowledgments

We thank Tatjana Lalic and Xinyu Li for providing technical support.

## Author contributions

Conceptualization: YW, LW, ZC

Data curation and analysis: LW, YW, FZ, GF, JVV, VW, AP, HS

Validation: LW, YW, GF, FZ

Methodology: LW, YW, FZ, GF

Investigation: LW, YW, FZ, GF, JVV, VW

Visualization: LW, FZ, ZC

Funding acquisition: ZC

Supervision: ZC, SP

Writing – original draft: ZC, LW

Writing – review & editing: ZC, LW

## Data and materials availability

All data are available on request.

## References

1. A. W. Fox, R. L. Davis, Migraine chronobiology. Headache 38, 436–441 (1998).

2. P. H. Newman, Median nerve compression in the carpal tunnel. Postgrad Med J 24, 264–269 (1948).

3. T. Tomson, K. Ekbom, Trigeminal neuralgia: time course of pain in relation to carbamazepine dosing. Cephalalgia 1, 91–97 (1981).

4. C. J. Glynn, J. W. Lloyd, The diurnal variation in perception of pain. Proc R Soc Med 69, 369–372 (1976).

5. R. S. Crockett, R. L. Bornschein, R. P. Smith, Diurnal variation in response to thermal stimulation: mouse-hotplate test. Physiol Behav 18, 193–196 (1977).

6. S. Farkas et al., Utility of different outcome measures for the nitroglycerin model of migraine in mice. J Pharmacol Toxicol Methods 77, 33–44 (2016).

7. G. A. Weir et al., The Role of TRESK in Discrete Sensory Neuron Populations and Somatosensory Processing. Front. Mol. Neurosci. 12, (2019).

8. P. Pettingill et al., A causal role for TRESK loss of function in migraine mechanisms. Brain 142, 3852–3867 (2019).

9. H. K. Kim et al., Circadian regulation of chemotherapy-induced peripheral neuropathic pain and the underlying transcriptomic landscape. Sci Rep 10, 13844 (2020).

10. G. T. van der Horst et al., Mammalian Cry1 and Cry2 are essential for maintenance of circadian rhythms. Nature 398, 627–630 (1999).

11. J. Lau et al., Temporal control of gene deletion in sensory ganglia using a tamoxifen-inducible Advillin-Cre-ERT2 recombinase mouse. Mol Pain 7, 100 (2011).

12. W. D. Todd et al., Suprachiasmatic VIP neurons are required for normal circadian rhythmicity and comprised of molecularly distinct subpopulations. Nat Commun 11, 4410 (2020).

13. T. Lalic et al., TRESK is a key regulator of nocturnal suprachiasmatic nucleus dynamics and light adaptive responses. Nat Commun 11, 4614 (2020).

14. T. Hirota et al., Identification of small molecule activators of cryptochrome. Science 337, 1094–1097 (2012).

15. S. Farkas, K. Bölcskei, A. Markovics, A. Varga, Á. Kis-Varga, V. Kormos, B. Gaszner, C. Horváth, B. Tuka, J. Tajti, Z. Helyes, Utility of different outcome measures for the nitroglycerin model of migraine in mice. J Pharmacol Toxicol Methods. 77, 33–44 (2016).

16. H. M. Jahn, C. V. Kasakow, A. Helfer, J. Michely, A. Verkhratsky, H. H. Maurer, A. Scheller, F. Kirchhoff, Refined protocols of tamoxifen injection for inducible DNA recombination in mouse astroglia. Scientific Reports. 8, 5913 (2018).

17. G. T. J. van der Horst, M. Muijtjens, K. Kobayashi, R. Takano, S. Kanno, M. Takao, J. de Wit, A. Verkerk, A. P. M. Eker, D. van Leenen, R. Buijs, D. Bootsma, J. H. J. Hoeijmakers, A. Yasui, Mammalian Cry1 and Cry2 are essential for maintenance of circadian rhythms. Nature. 398, 627–630 (1999).

18. S.-H. Yoo, S. Yamazaki, P. L. Lowrey, K. Shimomura, C. H. Ko, E. D. Buhr, S. M. Siepka, H.-K. Hong, W. J. Oh, O. J. Yoo, M. Menaker, J. S. Takahashi, PERIOD2::LUCIFERASE real-time reporting of circadian dynamics reveals persistent circadian oscillations in mouse peripheral tissues. Proc Natl Acad Sci U S A. 101, 5339–5346 (2004).

19. M. B. Elliott, M. L. Oshinsky, P. S. Amenta, O. O. Awe, J. I. Jallo, Nociceptive neuropeptide increases and periorbital allodynia in a model of traumatic brain injury. Headache. 52, 966–984 (2012).

20. S. R. Chaplan, F. W. Bach, J. W. Pogrel, J. M. Chung, T. L. Yaksh, Quantitative assessment of tactile allodynia in the rat paw. J. Neurosci. Methods. 53, 55–63 (1994).

21. W. J. Dixon, Efficient analysis of experimental observations. Annu. Rev. Pharmacol. Toxicol. 20, 441–462 (1980).

22. S. A. Malin, B. M. Davis, D. C. Molliver, Production of dissociated sensory neuron cultures and considerations for their use in studying neuronal function and plasticity. Nat Protoc. 2, 152–160 (2007).

23. A. Jagannath, R. Butler, S. I. H. Godinho, Y. Couch, L. A. Brown, S. R. Vasudevan, K. C. Flanagan, D. Anthony, G. C. Churchill, M. J. A. Wood, G. Steiner, M. Ebeling, M. Hossbach, J. G. Wettstein, G. E. Duffield, S. Gatti, M. W. Hankins, R. G. Foster, S. N. Peirson, The CRTC1-SIK1 pathway regulates entrainment of the circadian clock. Cell. 154, 1100–1111 (2013).

24. T. Lalic, A. Steponenaite, L. Wei, S. R. Vasudevan, A. Mathie, S. N. Peirson, G. S. Lall, M. Z. Cader, TRESK is a key regulator of nocturnal suprachiasmatic nucleus dynamics and light adaptive responses. Nat Commun. 11, 4614 (2020).

25. T. Dobler, A. Springauf, S. Tovornik, M. Weber, A. Schmitt, R. Sedlmeier, E. Wischmeyer, F. Döring-, TRESK two-pore-domain K+ channels constitute a significant component of background potassium currents in murine dorsal root ganglion neurones. J. Physiol. (Lond.). 585, 867–879 (2007).

26. P. Liu, Z. Xiao, F. Ren, Z. Guo, Z. Chen, H. Zhao, Y.-Q. Cao, Functional analysis of a migraine-associated TRESK K+ channel mutation. J. Neurosci. 33, 12810–12824 (2013).

