## Supplementary materials for "Cryptochrome Stabilization Ameliorates Chronic Pain"

Materials and Methods

Table S1-S4

Supplementary Fig 1-7

Reference 15-26

**Materials and Methods**

**Animals**

All procedures were carried out under an approved UK Home Office License, following the UK Home Office Animals (Scientific Procedures) Act 1986 and the University of Oxford’s Policy on the Use of Animals in Scientific Research. 2-3 month-old mice weighing 20-30g on a C57BL/6J background (Charles River) were housed in pathogen-free individually- ventilated cages with no more than 5 mice/cage. Mice were housed in 12:12-h light/dark (LD) cycles or constant dark (DD) conditions, for no more than 8 days, with food and water *ad libitum.* Transgenic mice were bred on a C57BL/6J background. This study conforms to the ARRIVE guidelines.

**Nitroglycerin (GTN) mouse model**

GTN (Hospira) was prepared for injection as 1mg/ml solution reconstituted in 0.9% NaCl. Following baseline behavioral testing, GTN (10mg/kg) or vehicle (0.9% NaCl) was administered (i.p.) 2h prior to testing (acute model) or alternative days for the chronic model. The experimenter was blind to both treatment and genotype; treatment was randomly assigned. Mice may be reused after 2 weeks of the previous test as all treatment has been eliminated from the body as previously described (*15*) but only reused once to maintain the moderate severity boundary and were consequently euthanized by schedule 1 culling techniques.

**GM mouse models**

*Advillin-Cre-ERT2:Bmal ^fl/fl^* mice: hemizygous *Advillin-cre-ERT2* tamoxifen-inducible mouse (JAX stock No. 032027) was crossed with homozygous *Bmal ^fl/fl^* mouse (JAX stock No. 007668) to produce hemizygous *Advillin-cre-ERT2* heterozygous *Bmal ^fl/fl^* that are crossed again with homozygous *Bmal ^fl/fl^* to obtain hemizygous *Advillin-cre-ERT2* and homozygous *Bmal ^fl/fl^*. Injection of tamoxifen (i.p. 100mg/kg dissolved in corn oil, 5 consecutive injections 24hr apart. Behaviour tests and sample collection 12-20 days from the first day of injection(*16*)) induced knock out of *Bmal1* in *Advillin-Cre* positive cells. Genotyping primers: *Advillin-cre-ERT2* wildtype band ~ 480bp forward: 5’-CCCTGTTCACTGTGAGTAGG; reverse: 5’-AGTATCTGGTAGGTGCTTCCAG. Mutant band ~ 180bp forward: 5’- CCCTGTTCACTGTGAGTAGG; reverse: 5’- GCGATCCCTGAACATGTCCATC. *Bmal ^fl/fl^* forward: 5’-ACTRRAARTAACTTTATCAAACTG; reverse: 5’-CTGACCAACTTGCTAACAATTA. Wildtype band ~327bp mutant band ~431bp.

*VIP-Cre:Bmal ^fl/fl^* mice: homozygous *VIP-Cre* mouse (JAX stock No. 031628) was mated with homozygous *Bmal ^fl/fl^* mouse (JAX stock No. 007668) to produce a heterozygous generation that was crossed again to obtain homozygous *VIP-Cre* and homozygous *Bmal ^fl/fl^* that results in *Bmal1* KO in *VIP-Cre* positive cells. Genotyping primers: *VIP-Cre* wildtype band ~175bp forward 5’-TCC TTG GAA CAT TCC TCA GC; reverse 5’-GGA CAC AGT AAG GGC ACA CA. mutant band ~350bp forward 5’-CCC CCT GAA CCT GAA ACA TA; reverse 5’-GGA CAC AGT AAG GGC ACA CA. *Bmal ^fl/fl^* forward 5’-ACTRRAARTAACTTTATCAAACTG; reverse 5’-CTGACCAACTTGCTAACAATTA. Wildtype band ~327bp mutant band ~431bp.

*Cry1*^-/-^:*Cry2*^-/-^ mouse: mouse colony with *Cry1* and *Cry2* knocked out sites as previously published(*17*). Genotyping primers: *Cry1* wildtype band ~499bp forward 5’- GGC CAT GTG GCT ATC TGT TC; reverse 5’- TAC AGG CAG CAG TGA GGA AG. *Cry2* wildtype band ~374bp forward 5’- AAG ACA GGC TTC CCT TGG AT; reverse 5’- GGA GCA GCT CGT CAA ATA CC. knocked out animal will have these bands missing.

*mPer2^Luc^* mouse: mouse colony with *mPer2^Luc^* knock-in that produced mPER2::LUC fusion protein as previously published(*18*). Genotyping primers: To distinguish between heterozygous and homozygous knock-in animals, PCR genotyping was performed by using 5’-CTGTGTTTACTGCGAGAGT-3 (P1) and 5’-GGGTCCATGTGATTAGAAAC-3 (P2) as WT allele detection primers and 5’-TAAAACCGGGAGGTAGATGAGA-3 (P3) as a reverse primer with P1 for the *Luc* knock-in allele detection.

**von Frey orofacial stimulation**

Mice were placed in a 15-30g head access restraining cylinder (Stoelting Co., Europe) and acclimatized for 5-10 minutes. The restraining cylinder allowed access to the head through various holes, with minimal ability for the mouse to move. Testing started when the mouse stopped exploring the cylinder (approximately 5 minutes) and a von Frey filament of 0.4g force was applied to the orofacial region at a 90° angle until the filament bent (*19*). The filament was applied 3 times and responses were recorded using a scoring system as follows: no response (0 point); mild response: gentle head shake/ forepaw swipe (0.25 points); moderate response: vigorous head shake/aggression to filament (0.50); and obvious response: complete withdrawal of the head/ facial grimacing (1 point). The points were summed to give a complete orofacial pain response score whereby a score of 3 is the highest response to record. An average was taken from left and right sides of the face. At least 48h was left between a battery of tests with the same animal to minimize variation in results.

**von Frey hind paw mechanical stimulation**

Static mechanical thresholds in freely-moving awake were examined via von Frey hair application (0.008–1 g; Touch Test; Stoelting) to the plantar surface of the hindpaw using the “up–down” method (*20*). Before testing, mice were randomly assigned to individual Plexiglas cubicles (8 x 5 x 10 cm) on an elevated wire mesh floor to enable access to the paw surface. Calibrated von Frey hairs were applied to the left and right hind paws, starting with the 0.6 g filament until the fiber bowed. A positive withdrawal response is followed by a lower force hair and vice versa for a negative withdrawal response until a behavior change occurs. Using this up–down sequence, four subsequent hairs were assessed and the 50% paw-withdrawal threshold (PWT) was calculated as described by Dixon (1980)(*21*). Data presented are from both left and right hind paws averaged together as no laterality effects were observed. At least 48h was left between a battery of tests with the same animal to minimize variation in results.

**Hargreaves test**

Thermal thresholds in freely-moving awake mice were assessed with the Hargreaves method using the plantar test (37370, UgoBasile). Before testing, mice were assigned to testing cubicles at random and habituated to the apparatus for 1 hour in individual cubicles (8 x 5 x 10 cm) placed on a glass plate. An infrared light source was applied to the plantar surface of hind-paws through the glass plate. Withdrawal reflexes were recorded from both left and right-paws, on three occasions, leaving at least 2 minutes between stimuli.

**Diurnal variation in nociception**

Wildtype mice were placed in 3 groups: Zeitgeber time (ZT)/ Circadian Time (CT) 1-3, 6-9 or 12-15 (n=12/group except for ZT6-9 n=21). Orofacial and von Frey hind paw stimulation testing was either completed in 12:12h LD cycle (ZT) or DD (CT). ZT=0 is when lights turn on in the animal facility; in DD mice were transferred to a completely dark chamber as soon as the lights were turned on which allowed an estimation of the circadian time as no wheel running or infrared beams were used to determine the exact activity/rest onset. Mice were left in constant darkness for 24h or 5 days and then throughout the testing period (3 days: training, baseline, testing). Dimmed red lighting was used during treatment administration and behavioral testing to allow the experimenter to see but little to no behavioral response change in mice due to different spectral sensitivity.

**TG and DRG primary culture**

Trigeminal ganglia (TG) and dorsal root ganglia (DRG) were dissected following a modified protocol from previously described by Malin et al. 2007 (*22*). Ganglia were collected in ice-cold Hanks' Balanced Salt Solution (HBSS). The DRGs of maximally two mice were extracted (±0.5 hour per mouse) in order to minimize time between dissection and plating. TG were quickly dissected and placed into HBSS.

TG were translocated to a 15ml tube and moved into a laminate flow hood for further processing. HBSS was carefully removed with a glass pipette and pre-warmed papain solution was added. Cells were incubated for 20min in a bead bath of 37 degrees. Tubes were placed at 45 degrees angle to increase the surface area of the tissue exposed to papain solution. Tapping of the tube was performed every few minutes to ensure proper contact with the solution. After 20 minutes, TG were centrifuged for 1 minute (<200g) and the papain solution was removed. Collagenase-dispase solution was added and TG were incubated for 20 minutes (repeating angular placement and tapping). After incubation, trituration was performed twice with a P1000 pipette. Cells were centrifuged for four minutes at 400g and Collagenase-dispase solution was removed. Pre-warmed dissociation media (DMEM/F12, FBS, P/S) containing DNAse (100ug/ml) was added. Trituration was performed three times with a P200 pipette and TG were incubated in the media containing DNAse at room temperature for 5 minutes. Trituration was then performed for twice with a P200 pipette. Pre-warmed dissociation media (=>3ml) was added to the cell suspension and then filtered through a 40um sterile sieve. The cell suspension was translocated to a 15ml tube and centrifuged for 6 min at 1000g. After removing the supernatant, the pellet was loosened, and the cells resuspended in 400ul dissociation media (+Rock inhibitor). Tissue culture wells were coated with PDL overnight (100ug/ml), washed with distilled water, and laminin was added (2ug/ml) and incubated for 1-6 hours. Cells were plated into two wells (200ul per well, 48-well plate); around 6.000-15.000 neurons per well. After two hours, the media was replaced with fresh growth media (Neurobasal, B27, Glutamax, P/S, NGF, GDNF). The media was changed the next day, and every two days after that.

**Bioluminescence recordings**

TG and SCN mouse explant culture

Mice were euthanized by cervical dislocation and a pair of trigeminal ganglia (n=4) and the suprachiasmatic nucleus (n=2) were rapidly dissected out of mPer2Luc knock-in mice from a C57BL/J6 background. Post- hoc statistical analysis suggested n=3 is the appropriate sample size for this current study (G-power, α= 0.05, power= 0.80). Individual ganglia were placed in chilled Hank’s Balanced Salt Solution (HBSS) and immediately transferred to millicell culture inserts (Merck Milipore PICM01250) in an opaque 24 well plate with recording media (Table 1) that contains reagents that do not interfere with circadian rhythms (*18*). Tissues were left to adjust to conditions in a 37^o^C incubator for approximately 10 minutes (*23*). Previous media was removed and recording media containing Luciferin (100uM, Gold Bio Technology; (*23*)) was added to each well. Plates were sealed on the bottom with opaque sealing film (Brightmax, Excel Scientific) and a transparent sealing film on top for accurate detection of bioluminescence without interference of other wells. Plates were placed in luminometer (Infinite 200 Pro, Tecan) at 36.5°C; each well recorded for 20 seconds every 30 minutes over approximately 5 days. Negative control containing only recording media and luciferin was used.

Circadian parameters (amplitude, period and phase) were calculated using Multicycle software (Acrimetrics). Baseline recordings can vary between different tissues, therefore data were detrended and smoothed into a polynomial curve, which is best-fit to a sine wave. There was no media change throughout the 5 days therefore tissue may be subject to decreased viability towards the end of the recording, which will show as a dampened amplitude, but it is important not to change media as this may affect the circadian analysis. Period analysis is the time it takes between peaks - one cycle is taken from 24h to 48h. Phase is made relative to one time point, 24h, in the SCN and is used to compare that to what phase the trigeminal ganglia is in at that time point.

Table S1. Recording Media

Reagents

DMEM Powder (Sigma D2902)
MiliQ water (1L)
Glucose (4.5ug/L, Thermofisher Scientific)
HEPES (1M, Thermofisher Scientific)
NaHCO3 (7.5%, Thermofisher Scientific)
Penicillin /Streptomycin (10,000U/ml, Thermofisher Scientific)

Glutamax (1X, Thermofisher Scientific)
B27 Plus Neuronal Culture Systems (1:50, Thermofisher Scientific)

Adjust pH to 7.3 and filter

DRG primary cell culture

DRG neurons were dissociated by primary cell culture protocol as described above, and plated on PDL and laminin coated 96 well plate with white walls (Greiner Bio-One Ltd #655983). After 5 days in culture, neurons were synchronized for an hour with either Forskolin 10uM or Dexamethasone 200nM against vehicle control (growth media: Neurobasal no phenol red, B27, Glutamax, P/S, NGF, GDNF). After an hour, the cells were washed once with PBS and then incubated with recording media (growth media + luciferin 100uM). Plates were sealed on the bottom with opaque sealing film (Brightmax, Excel Scientific) and a transparent sealing film on top for accurate detection of bioluminescence without interference of other wells. Plates were placed in luminometer (Infinite 200 Pro, Tecan) at 36.5°C; each well was recorded for 20 seconds every 30 minutes over approximately 2 days and a half. A negative control containing only recording media and luciferin was used.

**Neuron and SGC liquid-density-gradient dissociation**

This protocol is based primarily on Walker et al. (2014), using a gradient separation step to divide TG suspensions into two fractions: Larger neurons and glia (and some smaller neurons). The concentration of Dispase II is modified to comply with the manufacturer’s recommendation (0.6 - 2.4 U/ml).

P3-P6 mice are decapitated after being immobilized on ice. TGs are dissected out and collected in dissociation buffer on ice. After centrifuging at 100 x g for 3min to remove any contaminants in the supernatant, the TGs are allowed to digest and dissociate in a dissociation buffer containing Dispase II for 30min at 37 °C. Centrifuge at 200 x g for3 min and discard the supernatant. The pellet was resuspended in warm dissociation buffer, and triturated gently 15-25 times using a pipette (p1000). The suspension was centrifuge at 100 x g for 3 min and the pellet resuspend in L15 media. The layer gradient is prepared as: 6 ml L15+10 mg/mL BSA, carefully layered with 3 ml 1 mg/ml BSA on top without mixing before carefully layering the cell suspension onto the gradient. One final centrifuge at 100 x g for 3 min yields glia in the supernatant and neurons in the pellet.

Table S2. Reagents for cell fraction dissociation

| REAGENTS | Brand | Cat no |
| --- | --- | --- |
| DPBS without Ca^2^+ and Mg^2^+ | Gibco | 14190144 |
| HBSS without Ca^2^+ and Mg^2^+ | Gibco | 14170120 |
| HEPES | Sigma | H3375 |
| BSA | Sigma | A2153 |
| Dispase II | Gibco | 17105041 |
| L-15 Medium (Leibovitz) w/o L-glutamine | Sigma | L5520 |

**Immunocytochemistry (ICC)**

Cell cultures were rinsed once with PBS and then fixed with 4% paraformaldehyde (PFA) for 10 mins and rinsed twice with PBS. Cells were permeabilized with 0.1% Triton X-100 in PBS for 15 minutes. After removing Triton solution, cells were blocked with 10% donkey serum in PBS for 1 hour at room temperature. The blocked cells were incubated overnight with primary antibody in 5% donkey serum at 4^o^C, washed 3 times with PBS, and then incubated with secondary antibody in 5% donkey serum at room temperature for 1 hour. After washing 3 times with PBS and adding DAPI stain, the cells were mounted using either Dako mounting media (Agilent Technologies) or ProLong glass mounting media (ThermoFisher) before imaging with a EVOS fluorescence microscope. TG and SCN tissues were fixed with 4% PFA and then cryo-sectioned into 20-25um thick slices before ICC staining.

Table S3 Antibodies used for ICC

| Antibody | company | Used for | Dilution |
| --- | --- | --- | --- |
| Bmal1  Bmal1  TuJ1  Luciferase  IRDye680RD  IRDye800cw  Alexa 594 donkey anti-goat  Alexa 488 donkey anti-mouse  Alexa 647 goat anti-chicken | Abcam Ab235237  Cellsignal D2L7G  Sigma T8578-100ul  Abcam ab21176  Abcam ab216779  Abcam ab216774  Invitrogen  Invitrogen  Invitrogen | ICC  WB  ICC, WB  ICC  WB  WB  ICC  ICC  ICC | 1:200  1:1000  1:500; 1:1500  1:200  1:5000  1:5000  1:1000  1:1000  1:1000 |

**Western blotting**

TG tissues were rinsed in ice-cold PBS to remove excess blood and submerged in RIPA buffer with protease inhibitors. It was mashed up on ice with a homogeniser and sonicated for 10 mins at 4^o^C before centrifuged down (16000rpm for 20min at 4^o^C) to obtain supernatant in a fresh tube on ice and the pellet was discarded. Protein concentrations were determined with bicinchoninic acid assay (Pierce BCA assay kit, ThermoFisher Scientific 23227) at 562nm absorbance. NuPage system (Thermofisher) was used for running gels with MOPS SDS running buffer (NP00001) in mini gel tank with 200V for 30min. The nitrocellulose membrane midi from Bio-Rad (cat#1704159) was used with trans-blot turbo system for the protein transfer step. For blot visualization, the membrane was blocked in 7% donkey serum in PBS at room temperature for 2hr on a rocking platform or overnight at 4^o^C. The primary antibody solution was prepared in 4% donkey serum and the membrane incubated for 2 hours on a rocking platform or overnight at 4^o^C. The primary antibody solution was removed and the membrane was briefly washed with buffer (TBST/PBS +0.1% Twin20), followed by 10min buffer washes (x3). The secondary antibody solution was made with congregated IRDye in 4% blocking buffer and incubated with the membrane for 2 hours at room temperature on a rocker protected from light. After discarding the secondary antibody solution and washing the membrane 3 times, fluorescent imaging were obtained with the Odyssey fluorescent imaging platform.

**RT-qPCR**

All mice were euthanized via cervical dislocation and trigeminal ganglia were dissected and placed in RNAlater (Qiagen). RNA was extracted using Qiagen RNAeasy RNA extraction kit. Sufficient RNA quantity and purity for each sample were verified by RNA Nanodrop. RNA was reverse transcribed into cDNA using the Precision Nanoscript TM2 Reverse Transcription Kit from Primerdesign with random nanomer primers. Quantitative RT-PCR was performed using iTaq Universal probes supermix, probes from Taqman Thermofisher Scientific were used. All qRT-PCR samples were run with three technical repeats and the housekeeping gene B-actin served as the reference (GAPDH was also compared with B-actin, data not presented). All experiments were run on 7500 Fast (Applied Biosystems, Life Technologies) RT-qPCR machines using 7500 Software v2.3 (Life Technologies). Relative expression was completed using ΔCt method.

Table S4. Probe sequences (Taqman Gene Expression Assay, Thermofisher Scientific)

| Probe name | Probe |
| --- | --- |
| ActB  GAPDH | mm02619580_g1  mm99999915_g1 |
| TRESK | mm01702237_m1 |
| Trek1 | mm01323942_m1 |
| Trek2 | mm00504118_m1 |
| Per2  Per1  Bmal1  Cry1  Cyr2  Scn1a  Scn3a  Scn4a  Scn5a  Scn7a  SCN8a  SCN9a  Scn10a  Scn11a  Trpv1  GFAP  Trpa1  P2xr7  P2xr4  Kcna1  Nalcn  TrpM8  Nr1d1  Nr1d2 | mm00478099_m1  mm00501813_m1  mm00500226_m1  Mm01232937_m1  Mm01331542_m1  Mm00450580_m1  Mm07297464_m1  Mm00500103_m1  Mm01342518_m1  Mm00801952_m1  mm00488110_m1  mm00450762_s1  Mm00501467_m1  mm00449367_m1  mm01246300_m1  mm01253033_m1  mm01227437_m1  mm01199500_m1  mm00501787_m1  mm00439977_s1  Mm00616828_m1  Mm01299593_m1  Mm00520708_m1  Mm01310356_g1 |

**Calcium imaging**

Animals were decapitated after schedule 1 culling methods (isoflurane anesthesia or CO2). TG pairs were dissected out and blood vessels cleaned off as much as possible. TGs were immersed and positioned in 2% low setting-temp agarose in aCSF on a cutting plate immediately after dissection. The cutting plate was placed on ice to ensure fast gel formation, and then immersed in ice-cold cutting solution (oxygenated with 95% O2) on a vibratome. The gel block was trimmed before slicing with vibratome to minimize the cutting area. Vibratome settings: 210 - 320um thick slices at speed of 0.22mm/s. Slices were placed in oxygenated (5% CO2 and 95% oxygen) aCSF for about 10min before placing into Fura2 solution. A soft brush was used for transportation of slices between locations. Fura2-AM solution contained: 10uM Fura2-AM, 80uM pluronic acid, 1mg/ml collagenase 3, in aCSF. The Fura2-AM was incubated with pluronic acid for 2mins before adding into aCSF. Slices were kept in Fura2-AM solution for 1h at 25°C prior to recordings. The container was swirled gently to resuspend the slices every 15min to ensure that both sides of a slice had effective Fura2-AM penetration (5% CO2 and 95% oxygen while incubating). Slices were then washed and incubated in aCSF (5% CO2 and 95% oxygen) at room temp for at least 30mins before imaging. Baseline reading was taken with aCSF before adding ligands.

Cryoprotect ice-cold high-sucrose aCSF(*24*): 85 mM NaCl, 25 mM NaHCO3, 2.5 mM KCl, 1.25 mM 31 NaH2PO4, 0.5 mM CaCl2, 7 mM MgCl2, 10 mM glucose, and 75 mM sucrose

aCSF: 130 mM NaCl, 26 mM NaHCO3, 2.5 mM KCl, 1.25 mM NaH2PO4, 2 mM CaCl2, 1 mM MgCl2, and 10 mM glucose; pH7.4 Osmo 300

KCl solution: 100mM NaCl2, 100mM KCl2, 10mM HEPES, 10mM D-Glucose, 2mM Calcl2, 1mM MgCl2; pH7.4 with NaOH

**Chip-MEA electrophysiology**

TG slices were obtained the same way as described in the calcium imaging protocol. Slices placed in oxygenated (5% CO2 and 95% oxygen) aCSF for about 10min at room temperature before incubating in collagenase type 3 (1mg/ml; diluted in aCSF) for 1hr. The container was swirled gently to resuspend the slices every 15mins to ensure that both sides of the slice had effective penetration of collagenase. A single-chip MEA was used on the MEA2100-system (multichannel systems) to measure neuronal activity from TG slices. The well was filled with aCSF (infused with 5%CO2 and 95% O2) before placing a TG slice in the center, over the MEA electrodes. The slice was held down by a harp. Baseline recording was always taken before any stimulation.

**Patch clamp electrophysiology**

TNC slices

Whole-cell patch clamp recordings were obtained from brain slices from 6-8 weeks old male C57/BL6J mice. Brains, including the rostral portion of the cervical spinal cord, were extracted and submerged in an ice-cold cutting solution. The cutting solution was continuously bubbled with 95% O2/5% CO2 and contained (in mM): N-methyl D-gluconate (92), KCl (2.5), NaH2PO4 (1.25), NaHCO3 (30), HEPES (20), glucose (25), sodium ascorbate (5), MgSO4 (10), thiourea (2), sodium pyruvate (3), CaCl2 (0.5), osmolarity adjusted to 305-315 mOsm with sucrose. The pH was adjusted to 7.4 with HCl.

The cerebrum and cerebellum were removed and the remaining brainstem was glued on the side of the fourth ventricle onto the slicing table using cyanoacrylate. Horizontal slices of 250μm thickness containing the trigeminal nucleus were cut using a Leica VT1200S vibratome. Slices were immediately transferred to 34°C warm oxygenated NMDG for 10min. Slices were kept at room temperature in a custom-made holding chamber filled with aCSF for at least an hour before experimental use. aCSF was equilibrated to pH 7.3 to 7.4 when bubbled with 95% O2/5% CO2 and contained (in mM): NaCl (125), KCl (2.5), NaH2PO4 (1.25), NaHCO3 (26), glucose (10), CaCl2 (2.5), MgCl2 (1). The osmolarity was between 305 and 315 mOsm. Slices were kept for a maximum of 6 hours.

For recordings of spontaneous synaptic activity, slices were placed in an interface chamber and superfused with oxygenated aCSF at room temperature. Slices were visualized using a fixed-stage upright microscope fitted with a 40x water immersion objective. Cells were identified in the translucent area adjacent to the trigeminal tract using digital interference contrast. Recording electrodes were fabricated from borosilicate glass capillaries (World Precision Instruments, Hertfordshire, UK) using a Sutter P-1000 horizontal puller. Positive pressure was applied during the approach and released upon touching the cell surface, before switching to negative pressure to establish the giga-ohm (GΩ) seal. The plasma membrane was ruptured by the application of brief pulses of suction. Spontaneous excitatory postsynaptic currents (sEPSC) were recorded in voltage clamp configuration at a holding current of -70mV in presence of picrotoxin (30μM) using an internal solution containing (in mM): cesium chloride (130), HEPES (10), MgCl2 (6), Na2-ATP (2.5), Na-GTP (0.2), EGTA (4). The pH was adjusted to 7.3 with CsOH. Osmolarity was adjusted to 295 to 300mOsm with sucrose. At the end of each experiment, the glutamatergic origin of the recorded currents was confirmed by bath application of 2mM kynurenic acid. The liquid junction potential was not corrected. The amplifier head stage was connected to a MultiClamp 700B amplifier (Axon Instruments). The amplifier current output was filtered at 4 kHz and digitized at 20 kHz using an Axon Digidata 1550B (Axon Instruments). Data acquisition was performed using Clampex v10.6. Spontaneous EPSCs were detected using WinEDR (v3.8.6) using scaled template matching or threshold detection. Frequencies of sEPSCs were calculated from 30s time windows.

TG neurons from mouse primary culture

Primary mouse TG neurons were plated on ibidis dishes and cultured for up to 4 days in vitro before patch clamp experiment. Neurons were synchronized with Forskolin 10uM for an hour and experiments were performed in a fixed time frame at particular time window post synchronization. Cells were continuously superfused with oxygenated artificial cerebrospinal fluid aCSF (95% O_2_/5% CO_2_) containing 140 mM NaCl, 4.7 mM KCl, 2.5 mM CaCl2, 1.2 mM MgCl2, 10 mM HEPES and 10 mM glucose (pH7.3, 290-300 mOsm). Patch‐clamp electrodes (4–7 MΩ) were filled with an intracellular solution containing 130 mM KCl, 1 mM MgCl2, 5 mM MgATP, 10 mM HEPES and 0.5 mM EGTA; pH 7.3. Recordings were obtained using a Multiclamp 700B amplifier and digitized at 10–20 kHz using Digidata 1550 acquisition board and data was analyzed with Clampfit 10 (Molecular Devices). Cell capacitance was compensated for by the circuitry of the amplifier and series resistance was compensated 60-80% to reduce voltage errors. Upon establishing whole-cell access baseline biophysical properties such as cell capacitance, resting membrane resistance, and input resistance were measured and monitored throughout. Cells were abandoned if values exceeded -30mV.

For evaluation of potassium leak current, we used a previously reported ramp protocol (*25*, *26*). In voltage clamp, nociceptors were held at -60mV before depolarisation to -25mV for 300ms and then subsequently ramped to -135mV over 550ms. Outward currents were measured at the end of the 300ms step. Measurements were collected in the presence of 1uM TTX. Step protocols were run to check that action potentials were blocked before performing the ramp protocol.

For evaluation of voltage dependent sodium channel currents, the intracellular pipette solution contained 4.7 mM NaCl, 130 mM CsF, 0.5 mM EGTA, 1 mM MgCl2, 10 mM HEPES, 5mM MgATP, 10mM TEA-Cl. The pH was adjusted to pH 7.3 and 290 mOsm with 1 mM CsOH. In voltage clamp, nociceptors were maintained at a holding potential of -60mV, the membrane was hyperpolarised in a prepulse to -105mV for 90ms to remove inactivation of voltage gated sodium channels and then was depolarised to test potentials from -70mV to +20mV for 10ms in 10mV steps.

**Statistical analysis**

Data are represented as mean ± SEM. The number of animals in each data set is ≥3. Comparisons for statistical significance were assessed by one- or two-way ANOVA and post hoc multiple comparisons *t*-tests or unpaired *t*-tests using GraphPad Prism. Statistical analysis of calcium imaging results was done with a custom MATLAB (R2020b) script. Data from MEA was not normally distributed. This data, therefore, was log normalized before ANOVA analysis.

**Supplementary figures**

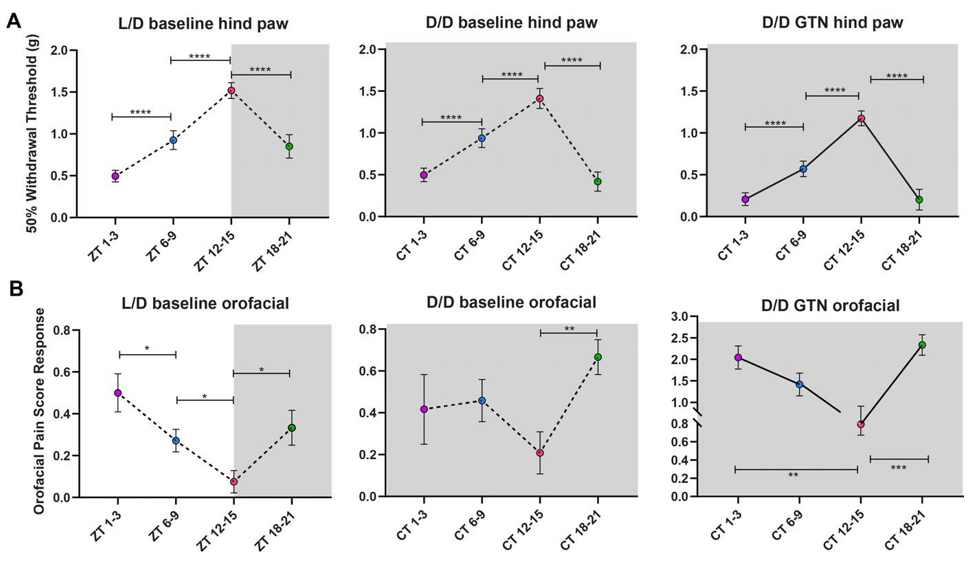

**Supplementary Fig. 1**

**(A**) Wildtype male 50% hindpaw withdrawal threshold at 4 time points in 12hr:12hr light/dark (L/D) condition (left), dark/dark (D/D) condition (middle) and GTN injected in D/D condition (right). n>=6 animals per time point. (**B**) wildtype male von Frey orofacial pain scores of the same animals from (A) of each condition. Graphs show mean +/- SEM; statistical significance with one-way ANOVA multiple comparisons test * p<0.05, ** p<0.005, *** p<0.0005, **** p<0.0001

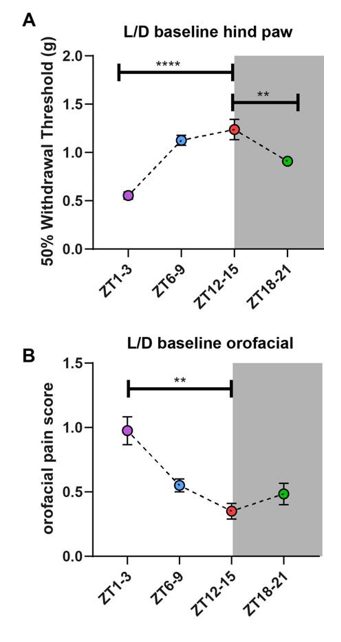

**Supplementary Fig. 2**

(**A**) Wildtype female 50% hindpaw withdrawal threshold at 4 time points in 12hr:12hr light/dark (L/D) condition (n=5 animals per time point). (**B**) wildtype female von Frey orofacial pain scores of the same animals from (**A**). Graphs show mean +/- SEM; statistical significance with one-way ANOVA multiple comparisons test * p<0.05, ** p<0.005, *** p<0.0005, **** p<0.0001

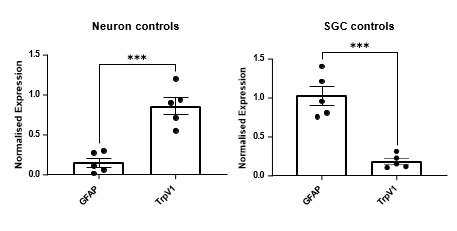

**Supplementary Fig. 3**

Gene expression comparison of *Gfap* and *Trpv1* in two groups (Neuron and SGC) after liquid density-based centrifuge separation (n=5 TG pairs/animals). Graphs show mean +/- SEM; statistical significance with two-tail unpaired *t* test *** p<0.0005.

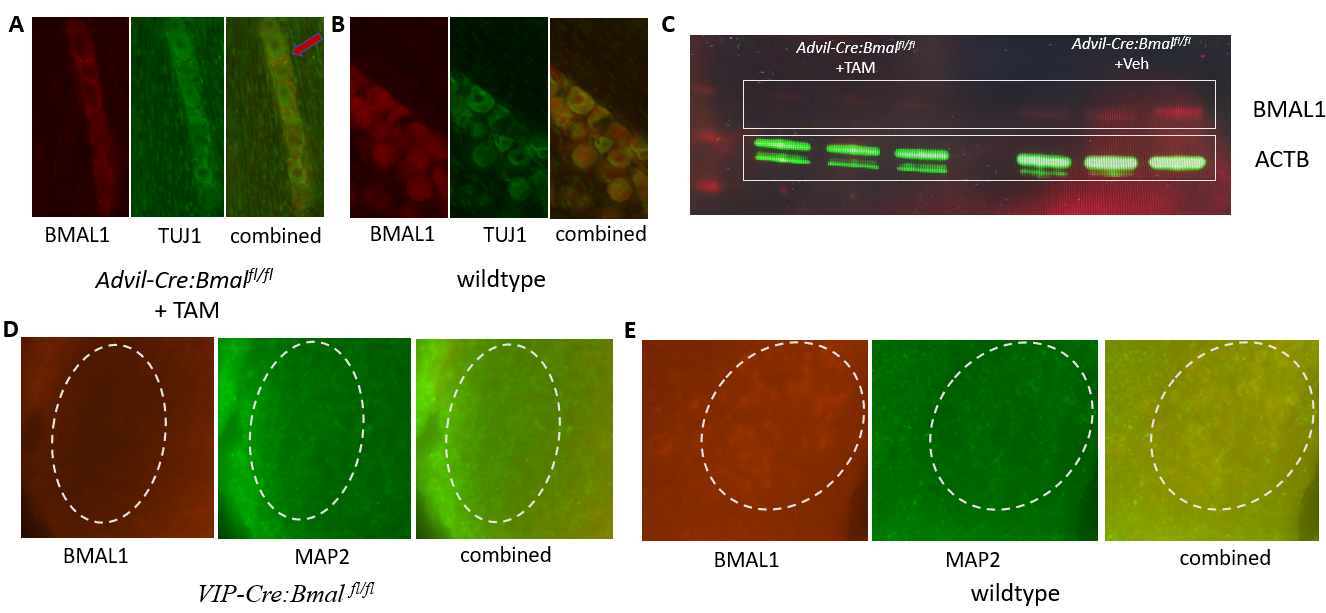

**Supplementary Fig. 4**

**(A**) Immunocytochemical staining for BMAL1 and TUJ1 in TG section of *Advillin-Cre-ERT2:Bmal ^fl/fl^* mice after i.p. Tamoxifen (TAM). The red arrow shows *Bmal1* in SGC around the TG neurons but not inside. (**B**) ICC staining of BMAL1 and TUJ1 in mouse TG of wildtype. BMAL1 is present inside neurons as well as SGCs. (**C**) Western blot of ACTB and BMAL1 for TAM induced *Bmal1* KO and control. Three TG per group. The double bands of ACTB are artificial from the blot transfer.

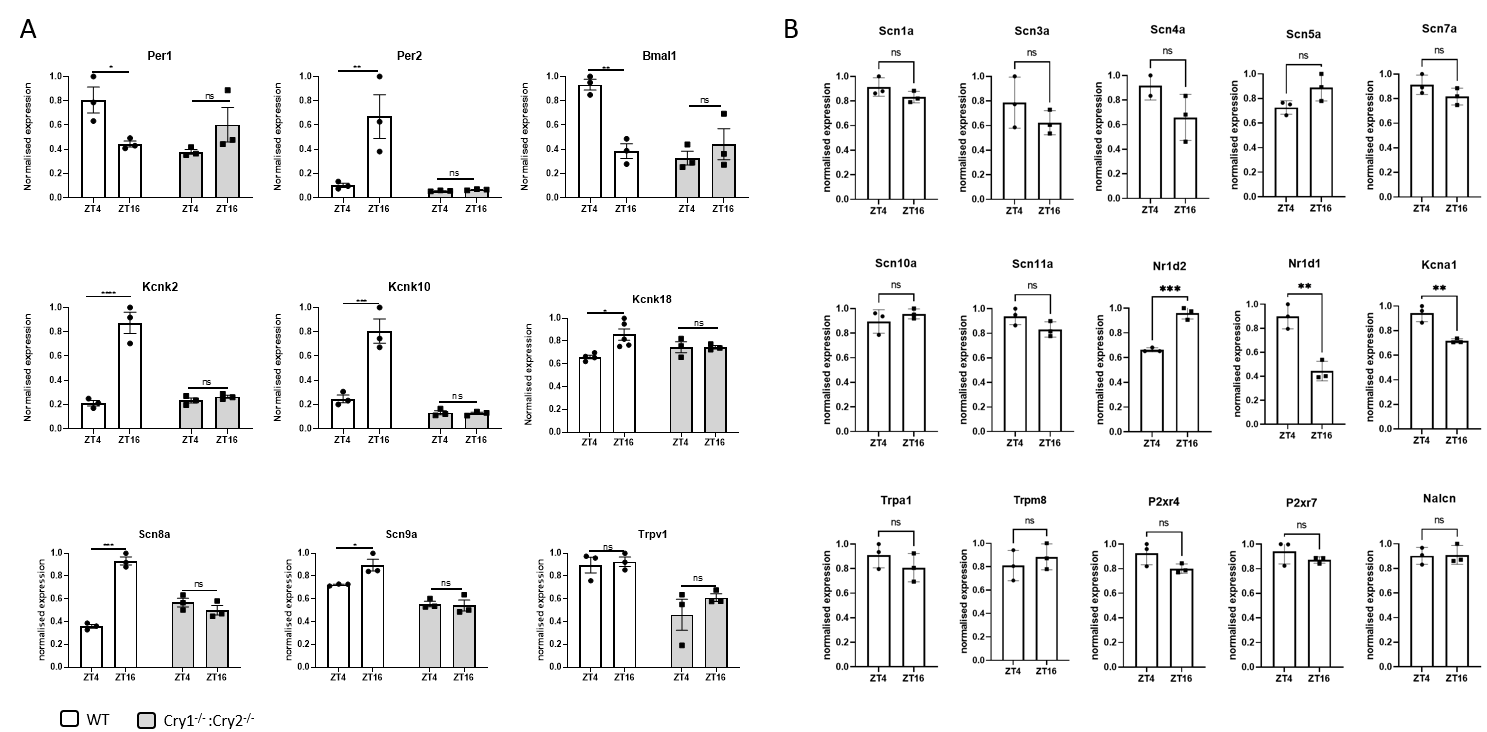

**Supplementary Fig. 5**

(**A**) Gene expression of clock genes and ion channel genes (*Per1*, *Per2*, *Bmal1*, *Kcnk2, Kcnk10, Kcnk18, Scn8a, Scn9a, Trpv1*) at ZT4 and ZT16 in wildtype (white bars) and *Cry1^-/-^:Cry2^-/-^*animals (grey bars). (**B**) Gene expression of ion channel genes (*Scn1a, Scn3a, Scn4a, Scn5a, Scn7a, Scn10a, Scn11a, Nr1d2, Nr1d1, Kcna1, Trpa1, Trpm8, P2xr4, P2xr7, Nalcn*) at ZT4 and ZT16 in wildtype animals. Graphs show mean +/- SEM of 3 biological repeats (3 technical repeats included) per time point. Statistical significance with two-tail unpaired t test * p<0.05, ** p<0.005, *** p<0.0005, **** p<0.0001.

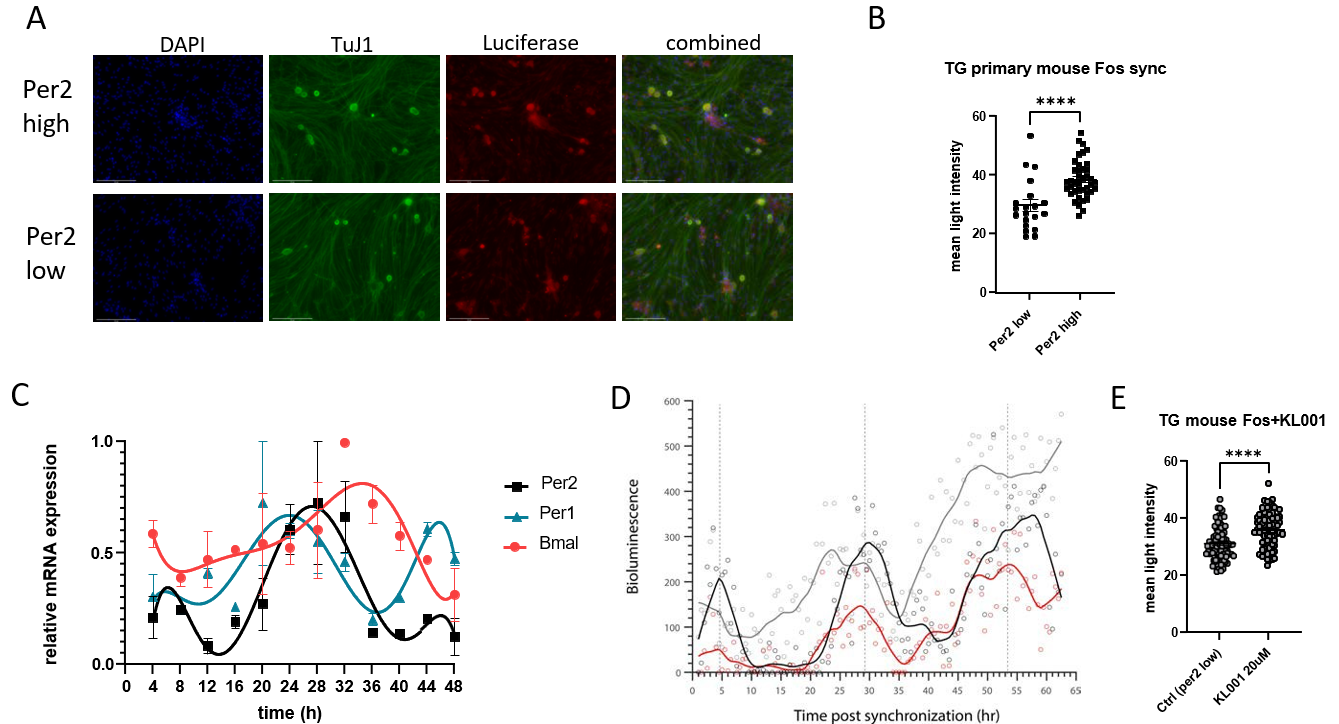

**Supplementary Fig. 6**

(**A**) ICC staining of *mPer2^Luciferase^* (*mPer2^Luc^*) knockin mice primary TG culture after Forskolin 10uM synchronization. Neurons were fixed and stained for TUJ1, LUC, and DAPI at *Per2* high (4hr post synchronization) and *Per2* low (16hr post synchronization). Light intensity images were taken at the same settings. (**B**) summary graphs of mean LUC light intensity per TG neuron at the two different time points after Forskolin synchronization (n>20 per time point). Statistical significance with two-tail unpaired *t* test **** p<0.0001. (**C**) relative clock gene expression (*Per2, Per1, Bmal1*) at 12 time points during 48hr post-Forskolin synchronization of mouse DRG primary neuron culture. Graphs show mean +/- SEM of 3 biological repeats (3 technical repeats included) per time point. (**D**) Bioluminescence data of mPER2::LUC in DRG primary neuron culture during 3 days post synchronization of Forskolin 10uM (black line), Dexamethasone 200nM (red line), and vehicle control (grey line). The lines in the graph are a smooth average of 3 biological repeats per condition. (**E**) summary graphs of mean luciferase light intensity per TG neuron at the 16hr post-Forskolin synchronization (*Per2* low time point) with vehicle or with KL001 in the media (n>20 per time point) Statistical significance with two-tail unpaired *t* test **** p<0.0001.

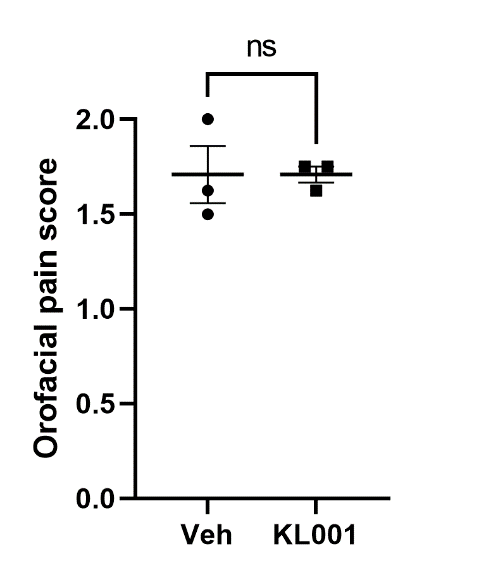

**Supplementary Fig. 7**

Orofacial pain score of *Cry1^-/-^:Cry2^-/-^* mice i.p. injected with either vehicle or KL001 (0.4mg/animal) at ZT0 and tested at ZT4. 3 male mice per condition. Statistical significance with two-tail unpaired *t* test.
